# Use of a citizen science tool for the determination of biological invasions in urban areas

**DOI:** 10.1101/2022.06.16.496492

**Authors:** Ana Luiza Moreira Botan, Gustavo Heringer, Ana Carolina Lacerda de Matos, Daniel Luiz Oliveira, Danielle Ramos de Alvarenga, Jonathan Wilson Almeida, Karla Palmieri Tavares, Marina Lopes Bueno, Vitor Hugo Lopes, Rafael Dudeque Zenni

## Abstract

Urban environments are critical points for biological invasions because these areas are susceptible to a greater number of environmental disturbances. Because they are densely populated ecosystems, urban environments present a unique opportunity for the involvement of society in the management of invasive exotic species. Similarly, citizen science offers opportunities to conduct research in the field of ecology together with society. The objective of our study was to analyze the occurrence of exotic species in urban areas recorded on iNaturalist and determine whether citizen science applications are good data sources for research projects in invasion ecology. Specifically, we evaluated whether richness and composition of the exotic species community in the cities registered on the iNaturalist platform were explained by socioeconomic and environmental factors. We also verified whether richness of the exotic species in cities registered on the iNaturalist platform were similar to the richness of the exotic species community in the region where the city is located using data collected only by researchers. We obtained 38,374 occurrences of 265 invasive species covering 2,057 cities in 72 countries. Occurrence records were concentrated in North America, Western Europe and Oceania, and there were no data for cities in most of Africa, the Middle East, Eastern Europe and West Asia. Socioeconomic and environmental factors were important determinants of the richness of exotic species in urban areas of the world and were important determinants of the richness of exotic species in natural environments. Urban hotspots of invasive exotic species were different from those for ecosystems in general.

## Introduction

The presence of invasive exotic species is commonly linked to the transport of goods and services and to human demography and migrations in tropical and subtropical regions (Essl et al. 2020). Urban environments in particular are critical points for biological invasions because these areas are susceptible to a greater number of environmental disturbances that can cause economic losses and harm the social well-being of human societies (Gaertner et al. 2016, Gaertner et al. 2017, Jackson 2015, Born et al. 2005).

Throughout the twentieth century, the urban human population grew from 20 million people to 2.8 billion, and today, more than half of the human population lives in urban areas (UNFPA, 2007). As a result, there is growing concern regarding habitat loss, the extinction of native species and the introduction of exotic species, as cities will continue to expand, change and fragment landscapes from agricultural and native areas to areas covered by impermeable surfaces (Aronson et al. 2017, Williams et al. 2015, Ricotta et al. 2017). The study of biological invasions has focused largely on the dissemination and impact of exotic species on natural and seminatural habitats. However, invasions are also widespread in human-dominated landscapes, where large economic costs usually occur due to altered ecosystem services, with impacts on human health and control efforts. Urban habitats represent the final pinnacle of the human modification of natural ecological and geological processes that structure a given location. Theories, concepts and paradigms for understanding and managing biological invasions are based largely on insights from natural and seminatural systems, but urban ecosystems differ radically from (semi)natural systems because the environmental impacts of human activity are harmful to many species; therefore, maintaining native biodiversity and ecosystem services that benefit people in cities can be difficult and expensive. Thus, it is necessary to monitor biological invasions in urban centers, and a tool that can assist in this monitoring is digital citizen science platforms.

Citizen science (voluntary involvement of people in the collection, analysis and interpretation of data) offers opportunities to conduct research together with society in the field of ecology (Adler et al. 2020, Pocock et al. 2017). Citizen science projects in ecology are able to cover a variety of subjects, involving millions of people around the world, contributing to an increase in the dissemination of science in human society (Bonney et al. 2014, Heigl et al. 2019). Digital citizen science platforms can be an option for obtaining useful data for studies on ecology and for monitoring invasive exotic species, in addition to potentially transmitting information to the population about the occurrence and consequences of biological invasions (Encarnação et al. 2021). An example of a digital platform that integrates researchers and people with enthusiasm for biology but without specific scientific training is iNaturalist. This application allows the identification of animals and plants through a photo taken and sent by the user.

Despite the great popularity and appeal of digital platforms utilized by the population to build large biodiversity databases, there is still some uncertainty about the quality and comprehensiveness of the data produced by people without specific scientific training (Haklay M. 2013). Thus, the objective of the present study was to analyze the occurrence of exotic species in urban areas recorded on the iNaturalist platform and determine whether citizen science applications can be used as reliable and sustainable data sources for research projects in invasion ecology. Specifically, we evaluated the richness and composition of the exotic species community in cities to determine the quantity and quality of exotic species data in urban areas, map the spatial distribution of the data, quantify and qualify the taxonomic diversity of the recorded data, list the most frequent and abundant exotic species in urban areas and analyze the feasibility of the application of iNaturalist in generating conclusions regarding urban biodiversity.

## Methodology

The data on the occurrence of exotic species were exported from the “Urban Exotic Biota” project, available on the iNaturalist platform (http://www.inaturalist.org; Accessed on September 21, 2020). There were no geographic or taxonomic restrictions in the search. To ensure that only observations in urban areas were considered, we overlapped the 192,091 observations obtained in iNaturalist with the presence of urban areas mapped using MODIS (version 4.0.0) data with a 0.5-km resolution (Schneider et al. 2009), and subsequently, we selected only the points of occurrence that overlapped with urban areas. The overlapping of the georeferenced observations from iNaturalist with the urban area map was performed in QGIS version 3.10.10. We also removed observations that were not considered research grade by the platform (research grade indicates data that are confirmed by more than two users). After applying these procedures, 83,074 observations remained for the analysis.

The exotic origin of the species in each urban area was verified using the Global Register of Introduced and Invasive Species (GRIIS - Pagad et al. 2018), available in the Global Biodiversity Information Facility (GBIF) network, and those observations of species that were not considered exotic in a given region by GRIIS were removed, leaving 38,374 observations. We used the GADM (Global Administrative Areas, 2012. GADM database of Global Administrative Areas, version 2.0. [online] URL: www.gadm.org.) database of global administrative areas as a classification standard for cities in the world. Thus, through geoprocessing tools, we associated the observations of exotic species obtained with city locations.

To determine which cities in the world had the greatest richness of exotic species recorded in iNaturalist, the biological groups observed by iNaturalist users and the environments where these observations were made, we calculated the total number of exotic species in each city and by biological group (Actinopterygii, Amphibia, Arachnida, Asteroidea, Aves, Bivalvia, Cephalaspidomorphi, Gastropoda, Hydrozoa, Insecta, Malacostraca, Mammalia, Maxillopoda, Polychaeta, Reptilia and Turbellaria) and by occurrence environment (freshwater, marine or terrestrial).

To study the composition of assemblages between regions, we performed a nonmetric multidimensional scaling (NMDS) ordination analysis using Jaccard distance. To avoid the bias caused by the high number of cities with few species occurrences, we restricted this analysis to cities with more than five species. We tested the clustering between regions with a permutational multivariate analysis of variance (“adonis” function, “vegan” package - Oksanen et al. 2021). The regions considered in this analysis are equivalent to the continents divided according to level 1 of the International Working Group on Taxonomic Databases (TDWG - Brummit 2001). Finally, we tested the correlation between gross domestic product per capita (GDP per capita), population density, human development index (HDI), mean annual temperature, annual rainfall and number of observations in iNaturalist and the clustering using the “envfit” function (“vegan” package - Oksanen et al. 2021). The GDP per country data were obtained from the World Bank for 2018 or the most recent year, and the HDI per country was obtained from the United Nations Development Program - 2020 Human Development Report. The mean annual temperature and annual rainfall data for each city were obtained from WorldClim 2 (Fick et al. 2017), and the number of observations per country was obtained from the iNaturalist platform (inaturalist.org) using the online platform iNaturalist Countries.

To identify the hotspots and coldspots of exotic species, we followed the same approach proposed by Dawson et al. (2017). As the distribution of the richness of exotic species can be influenced by the area of the city and the sampling effort (number of observations), we developed a linear model that took into account the effect of these variables. Next, we used the residual values to identify the cities with the highest and lowest richness of exotic species that were not explained by the model variables. The cities were divided into quantiles based on the richness (2.5%, 10% and 50% lower and 2.5%, 10% and 50% upper), where the cities with the highest occurrence of species (hotspots) were in the upper 2.5% quantile and cities with a lower occurrence of species (coldspots) in the lower 2.5% quantile. For the construction of the map, we used GADM data. Last, we evaluated the correspondence of the invasion hotspots and coldspots obtained with the data from the iNaturalist platform with the information provided by Dawson et al. (2017).

## Results

Of the 192,091 initial observations, we obtained 38,374 records of the occurrence of exotic invasive animals on seven continents, 72 countries and 2,057 cities (Fig. 1) for 265 different species. The occurrences of the species were distributed heterogeneously globally, with regions such as North America, Western Europe, Southeast Australia and New Zealand accounting for approximately 90% of the total records; there was a lack of information for large areas such as most of Africa, Saudi Arabia, Eastern Europe and Western Asia. The cities with the highest richness of exotic invasive animals were Toronto - Canada (n = 27), Auckland - New Zealand (n = 24) and Greater Vancouver - Canada (n = 23).

**Fig. 1.**
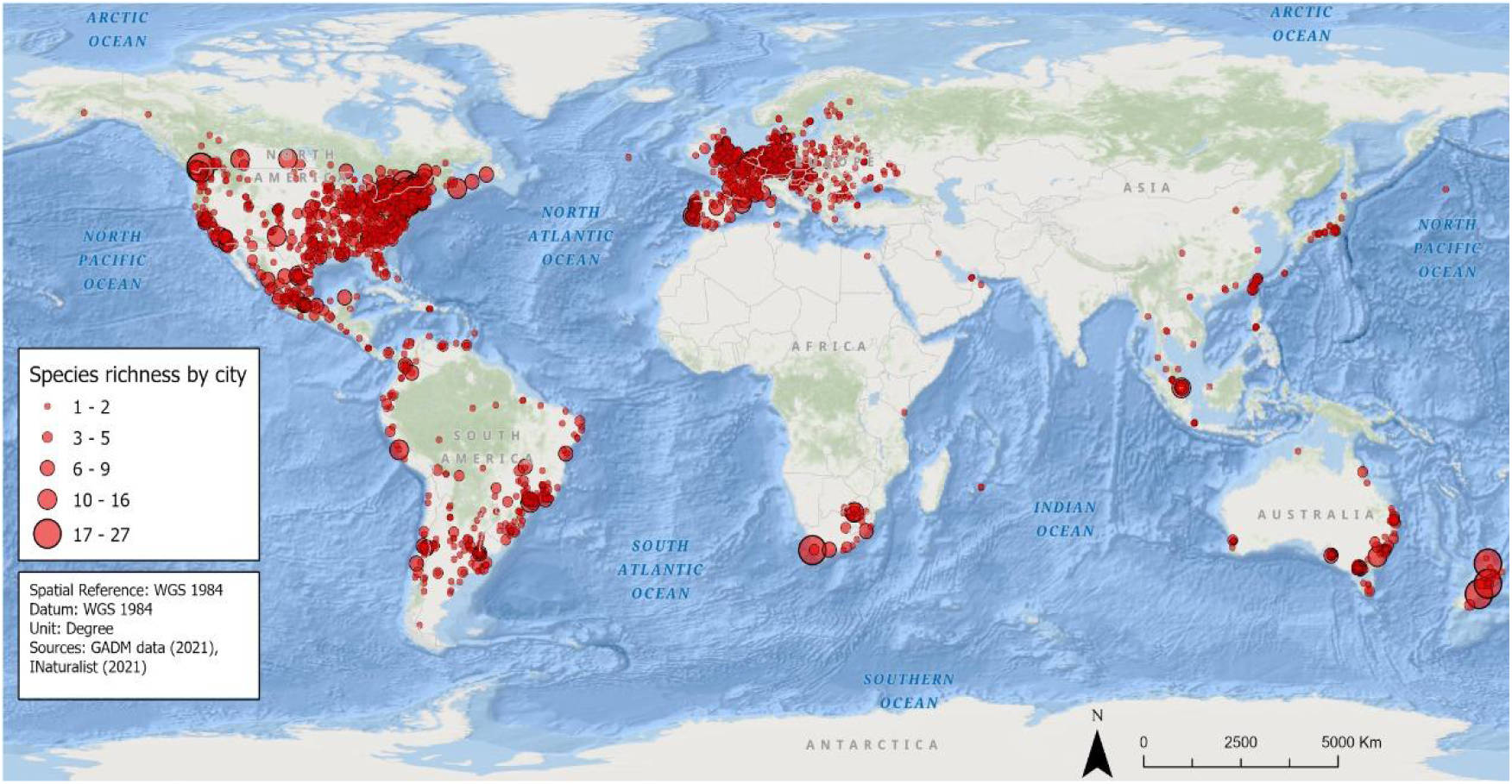
Richness of exotic species represented by city. Larger circles represent cities with a greater number of exotic species.

Among the taxonomic groups, Insecta had the highest species richness (n = 14 species), followed by Aves (n = 13 species) (Fig. 2). The other taxonomic groups showed richness lower than three species. For this analysis, we disregarded some taxonomic groups (Asteroidea, Cephalaspidomorphi, Hydrozoa and Polychaeta) because they had very low richness values. As seen in Figure 2, the cities with the highest insect richness were in Southern Africa, North America and Europe, and the cities with the highest Aves richness were in New Zealand, North America and South America.

**Fig. 2.**
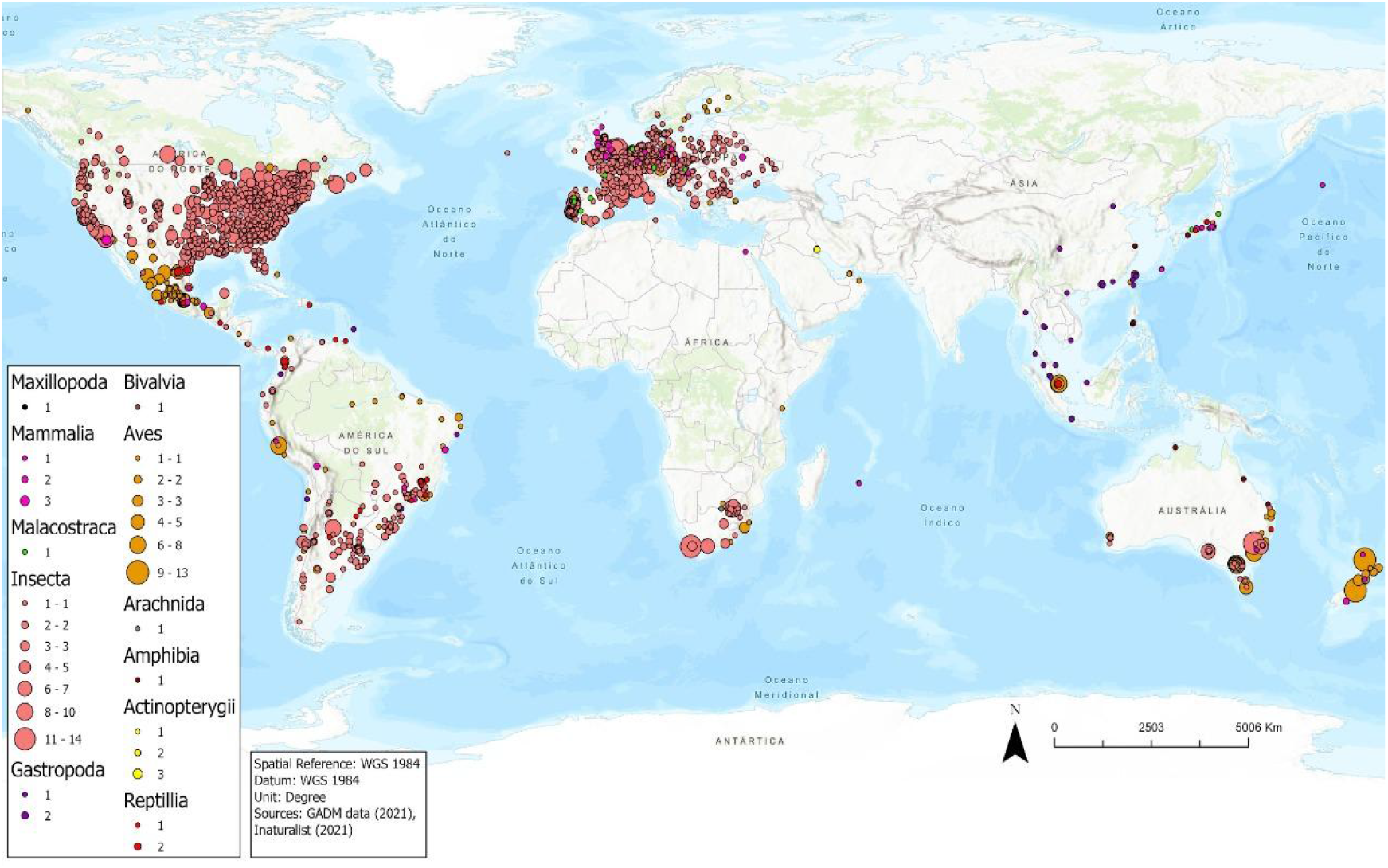
Invasive species richness represented by cities per taxonomic group. Larger circles represent cities with a greater number of exotic species.

Considering the three different environments (terrestrial, freshwater and marine) for each of the cities, we also found richness data per biological group per environment (Fig. 3). The only biological group present in all environments was Insecta, with a total richness of 85 species. For this group, the species that most appeared was *Harmonia axyridis*, with 24,560 observations, and it was also the species with the most observations among all species of all groups. In addition, the environment with the highest number of records was terrestrial (n = 236), followed by freshwater (n = 43) and finally marine (n = 10). Asteroidea, Cephalaspidomorphi, Hydrozoa, Maxillopoda and Polychaeta appeared in only one environment, and all of them had a richness equal to 1. Mammalia (n = 29), Actinopterygii (n = 21) and Arachnida (n = 19) presented richness higher than the previous groups but were present in only one environment.

**Fig. 3.**
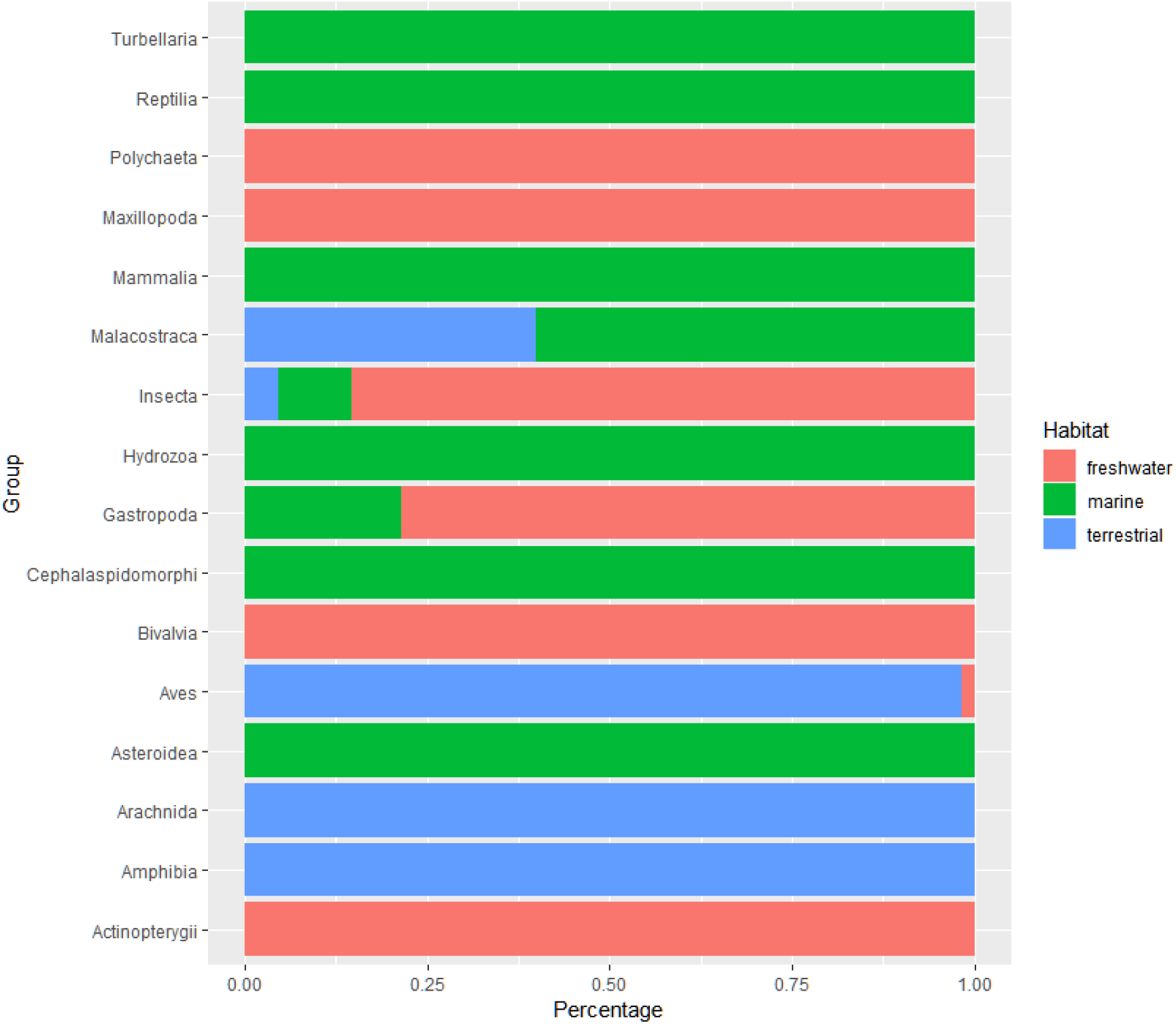
Percentage of species richness by biological group and city by environment. Orange bars represent freshwater species, blue bars represent terrestrial species and green bars represent marine species.

The composition of exotic invasive animals was affected by the continent where each city was located (PERMANOVA: p < 0.001). Cities located on the same continent showed species compositions more similar to each other. In addition, we found an association of environmental variables (i.e., rainfall and temperature), urbanization (i.e., gross domestic product per capita, population density, HDI and city area) and number of observations in iNaturalist with the ordination. The number of observations in iNaturalist, HDI and GDP were positively associated with the North American region, population density was positively associated with Europe, rainfall was positively associated with Eurasia and North America, and temperature were positively associated with tropical Asia and South America.

Through the analysis of hotspots, 28 cities had the highest residual values (upper 2.5% - Table 2) and 28 cities had the lowest residual values (lower 2.5%), corresponding to invasion hotspots and coldspots. The five main hotspot cities were Zama - Japan (residual = 14.61), Yokosuka - Japan (residual = 13.07), Timur Laut - Malaysia (residual = 10.57), Houten - Netherlands (residual = 9.64) and Hamm - Germany (residual = 8.59). The five main cold spot cities were Cachapoal - Chile (residual = −12.68), Muntinlupa - Philippines (residual = −8.64), Distrito Federal - Argentina (residual = −7.84), Panama - Panama (residual = −6.69) and Gaborone - Botswana (residual = −6.44).

**Table 1.** Data from the NMDS analysis. Permutations = 9999; stress = 0.11.

| Variables | NMDS1 | NMDS2 | R<sup>2</sup> | p |
| --- | --- | --- | --- | --- |
| Gross domestic product | -0.72948 | 0.68400 | 0.2318 | 0.0001 |
| Population density | 0.78134 | -0.62410 | 0.2134 | 0.0001 |
| Human development index | -0.94659 | 0.32243 | 0.1935 | 0.0001 |
| iNaturalist observations | -0.53699 | 0.84359 | 0.2902 | 0.0001 |
| City area | 0.88683 | -0.46210 | 0.0170 | 0.0016 |
| Rainfall | 0.78633 | 0.61781 | 0.0773 | 0.0001 |
| Temperature | 0.99941 | -0.03448 | 0.2301 | 0.0001 |

**Table 2.** City hotspots for the occurrence of exotic species and their respective residual and urban percentile values compared regional percentile from Dawson et al. (2017)

| City | Country | Residual | Urban percentile | Regional percentile |
| --- | --- | --- | --- | --- |
| Vlaams Brabant | Belgium | 4.16 | Upper 2.5% | Upper 50% |
| Soledad de Graciano Sánchez | Mexico | 4.16 | Upper 2.5% | Lower 50% |
| Hagen | Germany | 4.23 | Upper 2.5% | Upper 50% |
| Hume | Australia | 4.42 | Upper 2.5% | Upper 50% |
| Iztapalapa | Mexico | 4.53 | Upper 2.5% | Lower 50% |
| Brimbank | Australia | 4.72 | Upper 2.5% | Upper 50% |
| San José | Costa Rica | 4.73 | Upper 2.5% | Upper 50% |
| Kerkrade | Netherlands | 4.77 | Upper 2.5% | Upper 50% |
| Mayen-Koblenz | Germany | 4.86 | Upper 2.5% | Upper 50% |
| Eindhoven | Netherlands | 4.93 | Upper 2.5% | Upper 50% |
| Wien | Austria | 5.08 | Upper 2.5% | Upper 50% |
| Coacalco de Berriozábal | Mexico | 5.14 | Upper 2.5% | Lower 50% |
| Rio de Janeiro | Brazil | 5.22 | Upper 2.5% | Upper 50% |
| Nezahualcóyotl | Mexico | 5.80 | Upper 2.5% | Lower 50% |
| Camaçari | Brazil | 5.81 | Upper 2.5% | Upper 50% |
| Duiven | Netherlands | 5.82 | Upper 2.5% | Upper 50% |
| Ludwigshafen am Rhein | Germany | 5.84 | Upper 2.5% | Upper 50% |
| Campinas | Brazil | 6.44 | Upper 2.5% | Upper 10% |
| Praha 1 | Czech Republic | 6.58 | Upper 2.5% | Upper 50% |
| Liège | Belgium | 6.71 | Upper 2.5% | Upper 50% |
| Bruxelles | Belgium | 7.23 | Upper 2.5% | Upper 50% |
| Emmen | Netherlands | 7.79 | Upper 2.5% | Upper 50% |
| Unincorporated<br>Australian Capital<br>Territory | Australia | 7.89 | Upper 2.5% | Lower 50% |
| Hamm | Germany | 8.59 | Upper 2.5% | Upper 50% |
| Houten | Netherlands | 9.64 | Upper 2.5% | Upper 50% |
| Timur Laut | Malaysia | 10.57 | Upper 2.5% | Upper 50% |
| Yokosuka | Japan | 13.07 | Upper 2.5% | Upper 10% |
| Zama | Japan | 14.61 | Upper 2.5% | Upper 10% |

## Discussion

The iNaturalist platform proved to be a rich source of information on the occurrence of exotic species in urban areas, with data available for cities on all continents, except Antarctica. There was a predominant volume of information for North America, Western Europe and Oceania, but there were many records for cities in South America, South Africa and Asia (Fig. 1). When using only the occurrences recorded as being research grade, it was still possible to obtain an appropriate amount of data for macroscale comparative analyses (more than 80,000 observations). iNaturalist platform provides data in sufficient quantity and quality for research projects in invasion ecology that require information on the occurrence of exotic and invasive exotic species, provided that the existing biases in the data are considered in the analyses. An example of this is the paper developed by Ceccolini (2021), which used observations of the iNaturalist platform to identify the presence of the wasp *Sceliphron curvatum* in new locations. With the data obtained, it was possible to point out the first records of the species in new Portuguese cities, data that can be used to prevent new invasions of the species.

The distribution of richness of exotic species in urban areas of the planet depended on the same variables that explain the richness distribution in natural ecosystems. Socioeconomic factors (GDP, HDI and population density) and environmental factors (area, rainfall and temperature) are important determinants of the richness of exotic species in urban areas of the world as well as important determinants of the richness of exotic species in natural environments (Guimarães Silva et al. 2020, Van Kleunen et al. 2015, Zenni 2015). It is not surprising that factors that determine species richness in urban and natural areas are similar, but this finding reinforces the robustness of the data available in iNaturalist for surveying exotic urban biodiversity.

Based on the data available in iNaturalist, the urban hotspots of exotic species are different from those for ecosystems in general. When we compared the result of the hotspot analysis for the iNaturalist data with the work performed by Dawson et al. (2017), we observed that there was no correspondence between the hotspot and coldspot sites (upper and lower 2.5%). Among the 28 cities with the lowest residual values, only six corresponded to the lower percentiles (lower 10% and 50%) for the regions. Most hotspots corresponded to the upper percentiles found by Dawson et al. (2017), but none of them were associated with the regions with a greater distribution of invasive species (Table 2).

An important limitation observed in the iNaturalist data is that locations with more observations of invasive exotic species are also those with higher numbers of users on the platform. We know that the greater the number of people accessing the application, the greater the amount of available information, generating a possible observation bias. For example, we found that the terrestrial environment had the most observations, which is probably due to the ease of access by users. In addition, Insecta had, according to the taxonomic group, the highest number of observations and, consequently, the highest species richness in the platform. Aves closely followed Insecta, a pattern that may explain the fact that both are the taxonomic groups with the highest species richness.

A second limitation observed in the platform data is that more charismatic and visually attractive species also receive more attention from platform users. For example, *Harmonia axyridis*, a ladybug widely found in gardens and backyards and a common character in children’s stories, comics and cartoons, is the third insect and the fourth species with the most observations on the platform and a frequent species in our data. Data for plants, notoriously difficult to identify only by photography, were rarely of research grade and therefore were underrepresented in our dataset, as were fungi and microorganisms in general.

In conclusion, our results suggest that the iNaturalist platform offers robust and coherent data for the generation of conclusions regarding exotic urban biodiversity provided that the existing spatial and taxonomic biases are considered. Although the centers of exotic species richness in cities have diverged from the regional centers of exotic species richness, the same factors that determine the richness in natural environments have been shown to be important for urban environments. These results allow us to speculate that the exotic species existing in cities are not necessarily a subsample of the regional set of species. The incentive to use the iNaturalist platform by more people, especially in regions where the application has few users, and the incentive for taxonomists to participate in the platform to confirm the identification of species may greatly improve the availability of data in the future.

## Acknowledgments

ALMB thanks UFLA/CNPq for the scientific initiation scholarship. GH, ACLM, DRA, JWA and MLB thank Capes for the graduate scholarships. RDZ received partial financial support from CNPq (process 304701/2019-0).

## References

Adler, F. R., Green, M. A. & Sekercioglu, Ç. H. (2020). Citizen science in ecology: a place for humans in nature. Annals of the New York Academy of Sciences, 1–13.

Bonney, R., Shirk, J. L., Phillips, T. B., et al. (2014). Next Steps for Citizen Science. Science, 343, 1436–1437.

Born, W., Rauschmayer, F. & Bräuer, I. (2005). Economic evaluation of biological invasions – a survey. Ecological Economics, 55, 321–336.

Ceccolini, F. (2021) New records for the alien mud-dauber wasp *Sceliphron caementarium* (Drury, 1773) (Hymenoptera: Sphecidae) in Peru. Revista Chilena De Entomología, 47(4), 951–954.

Dawson, W., Moser, D., van Kleunen, M., et al. (2017). Global hotspots and correlates of alien species richness across taxonomic groups. Nature Ecology & Evolution, 1(0186).

Encarnação, J., Teodósio, M. A. & Morais, P. (2021). Citizen Science and Biological Invasions: A Review. Frontier in Environmental Science, 8.

Essl, F., Lenzner, B., Bacher, S., et al. (2020). Drivers of future alien species impacts: An expert-based assessment. Global Change Biology, 00, 1–14.

Fridley, J. D., Rebecca, L. B. & John, F. B. (2004). Null models of of exotic invasion and scale-dependent patterns of native and exotic species richness. Ecology, 85(12), 3215–3222.

Gaertner, M., Larson, B. M. H., Irlich, U. M., et al. (2016). Managing invasive species in cities: A framework from Cape Town, South Africa. Landscape and Urban Planning, 151, 1–9.

Gaertner, M., Wilson, J. R. U., Cadotte, M. W., et al. (2017). Non-native specien in urban environments: patterns, processes, impacts and challenges. Biological Invasions, 19, 3461–3469.

Global Administrative Areas. (2021). GADM Database of Global Administrative Areas. Accessed (xx). http://www.gadm.org/

Guimarães Silva, R., Zenni, R. D., Rosse, V. P., Bastos, L. S., & van den Berg, E. (2020). Landscape-level determinants of the spread and impact of invasive grasses in protected areas. Biological Invasions, 22(10), 3083–3099. https://doi.org/10.1007/s10530-020-02307-4

Güneralp, B., Lwasa, S., Msundire, H., et al. (2017). Urbanization in Africa: challenges and opportunities for conservation. Environmental Research Letterts, 13(1), 1–8.

Jackson, T. (2015). Addressing the economic costs of invasive alien species: some methodological and empirical issues. International Journal of Sustainable Society, 7(3), 221–240.

Haklay, M. (2013). Citizen Science and Volunteered Geographic Information – overview and typology of participation. In: Sui, D.Z., Elwood, S. and M.F. Goodchild (eds.), 2013. Crowdsourcing Geographic Knowledge: Volunteered Geographic Information (VGI) in Theory and Practice, 105–122.

Heigl, F., Kieslinger, B., Paul, K. T., et al. (2019). Toward na international definition of citizen science. PNAS, 116(17), 8089–8092.

Hochmair. H. H., Scheffrahn, R. H., Basille, M., et al. (2020). Evaluating the data quality of iNaturalist térmite records. PLoS ONE, (15)5.

Macic, V., Albano, P. G., Almpanidou, V., et al. (2018). Biological Invasions in Conservation Planning: A Global Systematic Review. Frontiers in Marine Science, 5.

Pocock, M. J. O., Tweddle, J. C., Savage, J., et al. (2017). The diversity and evolution of ecological and environmental citizen science. PLoS ONE, 12(4).

Schneider, A., M. A. Friedl and D. Potere (2009) A new map of global urban extent from MODIS data. Environmental Research Letters, volume 4, article 044003.

van Kleunen, M., Dawson, W., Essl, F., Pergl, J., Winter, M., Weber, E., Kreft, H., Weigelt, P., Kartesz, J., Nishino, M., Antonova, L. A., Barcelona, J. F., Cabezas, F. J., Cárdenas, D., Cárdenas-Toro, J., Castaño, N., Chacón, E., Chatelain, C., Ebel, A. L., … Pyšek, P. (2015). Global exchange and accumulation of non-native plants. Nature, 525(7567), 100–103. https://doi.org/10.1038/nature14910

Zenni, R. D. (2015). The naturalized flora of Brazil: A step towards identifying future invasive non-native species. Rodriguésia, 66(4), 1137–1144. https://doi.org/10.1590/2175-7860201566413

